# Nanomotion technology for testing azithromycin susceptibility of *Salmonella enterica*

**DOI:** 10.1101/2024.09.05.611511

**Authors:** Mariliis Hinnu, Toomas Mets, Ivana Kerkez, Marta Putrinš, Niilo Kaldalu, Gino Cathomen, Marta Pla Verge, Danuta Cichocka, Alexander Sturm, Tanel Tenson, ERADIAMR consortium

## Abstract

Azithromycin is used to treat invasive salmonellosis, despite conflicting effective concentrations *in vitro* and *in vivo*. Resistance of *Salmonella enterica* to azithromycin is increasing. We demonstrate that nanomotion technology can be used for rapid phenotypic testing of *Salmonella*’s susceptibility to azithromycin. Nanomotion changes under various culture conditions correlated with susceptibility measured by MIC determination, CFU counting, and fluorescent reporter-based estimates of intrabacterial azithromycin accumulation.

## MAIN

Invasive salmonellosis caused by *Salmonella enterica* subspecies is a major threat to human health affecting >20 million people yearly (1, 2). Antimicrobial resistance to traditional drugs, such as beta-lactams and fluoroquinolones, has emerged in all invasive *S. enterica* serovars (1). The macrolide azithromycin (AZI) has been effectively used to treat *Salmonella* infections resistant to other drug classes (3–5). AZI remains effective *in vivo*, despite recommended doses achieving peak serum concentrations in the range of 0.4 μg/ml (6): 20-fold lower than the MICs for most clinical strains (8 μg/ml) (7). Resistance to AZI is increasing (8, 9), underlining the need for rapid susceptibility testing. Nanomotion technology can be used as a rapid phenotypic antimicrobial susceptibility test (AST) (10–14). The technology is based on measuring oscillations caused by metabolically active organisms attached to a nanomechanical sensor, a cantilever (11, 15). The classification into resistant/susceptible categories is based on machine learning algorithms for specific strain-drug combinations. The susceptibility phenotype can already be detected two hours after blood culture positivity (14). The technology has been successfully applied in various bacterial species, and two clinical studies have been concluded (NANO-RAST (16), NCT05002413) and PHENOTECH- 1 (14), NCT05613322).

Prior to this study, nanomotion had not been used to determine susceptibility to AZI or any other macrolide. We recorded nanomotion of *S. enterica* under various experimental conditions affecting its susceptibility to AZI. We used neutral and acidic media, and two different incubation temperatures. In the early stages of development, nanomotion was measured at ambient room temperature (RT). The current setup uses 37°C for all ASTs to mimic physiological conditions in humans and to decrease the time to results (14).

Based on MIC values, *Salmonella* is up to 4 times more sensitive to AZI at RT compared to 37°C in different growth media (Fig. S1A; Table S1). This effect cannot be fully explained by the differences in growth rates (Fig. S1B). Nanomotion was recorded for AZI-susceptible *S. enterica* serovar Typhimurium SL1344 (17–19) (wild-type; wt) during AZI treatment and subsequent recovery in fresh drug-free medium at both RT and 37°C (Fig. 1 & S2). Before the addition of the antibiotic, nanomotion variance over time increased, indicating the presence and physiological activity of live bacteria on the cantilever. In the untreated sample, the signal continued to increase during the measurement (Fig. 1A). When AZI was added at concentrations exceeding the MIC, the nanomotion signal slope decreased. After removal of AZI nanomotion started to increase again in fresh drug-free medium at 37°C, indicating recovery (Fig. 1C; S2). However, no recovery was observed when the experiments were conducted at RT (Fig. 1B; S2), except after treatment with 16 μg/ml AZI (Fig. S2). In all cases, bacteria remained on the cantilever at the end of the experiment (Fig. S3).

**Figure 1.**
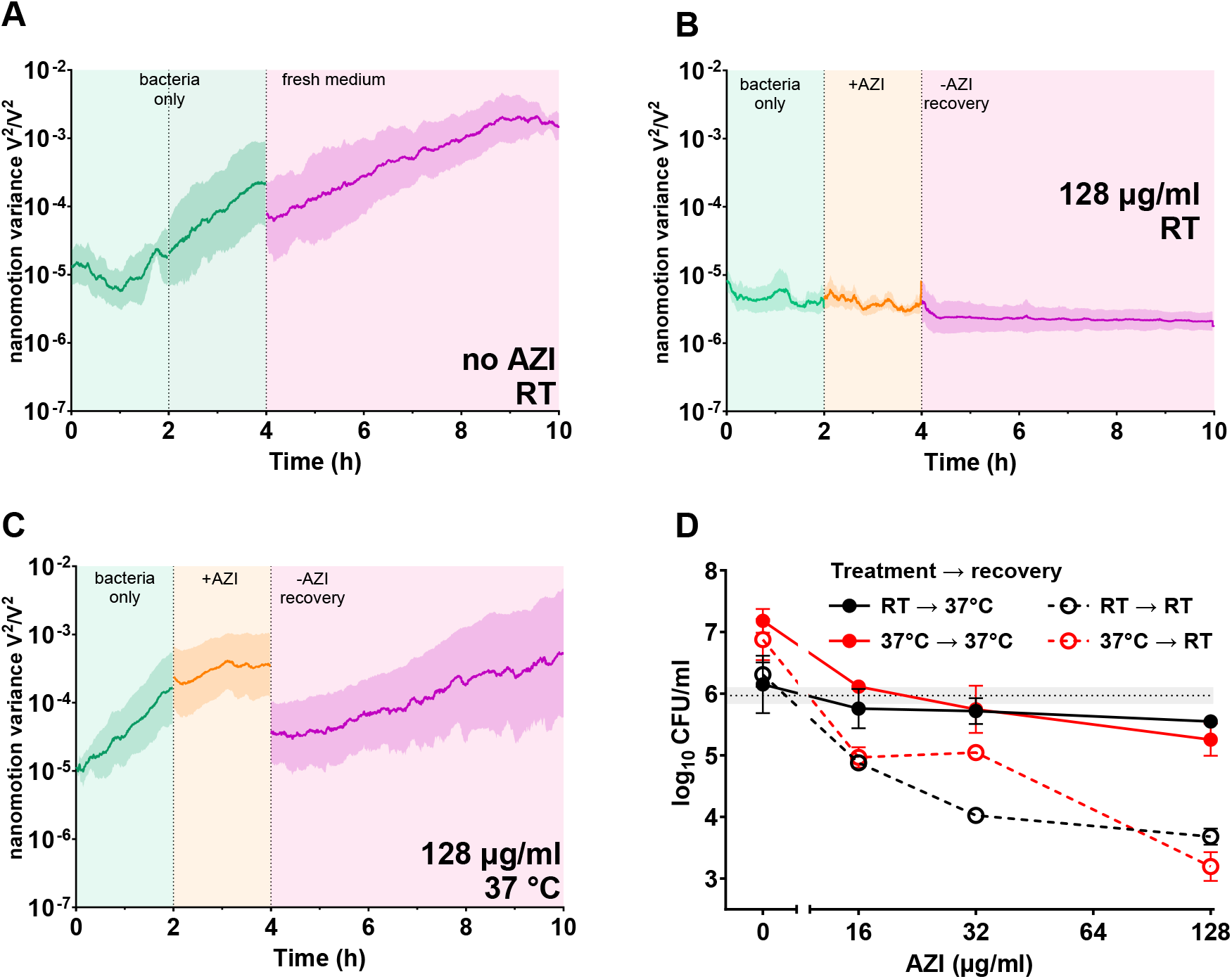
Variance over time of the nanomotion signal measurements of wt *Salmonella* without AZI (A) or with 128 µg/ml AZI for 2 hours and subsequent recovery in fresh medium at room temperature (RT) (B) or at 37°C (C). Green indicates bacterial nanomotion variance signal before adding the drug, orange is during drug treatment and pink is the recovery in fresh medium after removing the drug. Means ± SEM (N ≥ 3 biological replicates) shown for nanomotion data. D. Recovery of wt *Salmonella* colonies on LB-agar after 2h of treatment with AZI at indicated temperature at pH 7.4. Grey dotted line indicates the initial inoculum. Means ± SD (N ≥ 3 biological replicates).

We hypothesized that the bacteria might have been killed or their recovery delayed. Delay in post-treatment recovery after an antibiotic is removed from the extracellular environment is known as the postantibiotic effect (PAE), and it impacts antibiotic dosing (20, 21). Colony counts after treatment indicated that AZI killed less than one log of *S. enterica* when plates were incubated at 37°C during recovery. However, during RT recovery, the same concentrations of AZI killed at least one log more irrespective of the treatment temperature (Fig. 1D). The enhanced post-treatment killing by AZI at a lower temperature may reflect slower dissociation of the drug from the ribosome, which is known to increase the bactericidal activity of macrolides (22).

The slope of the nanomotion variance during drug exposure is a proxy for estimating drug susceptibility (14). To test whether detection of AZI resistance in *Salmonella* is feasible with nanomotion, we determined the slope of the variance at different AZI concentrations for wt SL1344 and a resistant mutant *acrB* R717Q, which harbors a clinically relevant mutation that increases AZI efflux in the acrAB-TolC efflux pump and has an MIC of 32 μg/ml (8, 23–25). We also determined the slope of the variance for wt strain at acidic pH, a condition encountered by intracellular *Salmonella* in acidic vacuoles (26) which increases AZI’s MIC above 1024 μg/ml (Table S1) (18). We used the rolling regression method for slope estimation, which demonstrated better reliability and robustness compared to the methods employed in previous studies (Supplementary materials and methods, Figure S4).

Remarkable differences in nanomotion arise between the strains at AZI concentrations near the MIC value of the wt at neutral pH. The drug slope values of the resistant mutant begin decreasing at higher AZI concentrations than the wt (Fig. 2A). A comparable difference is seen in the wt between neutral and acidic pH (Fig. 2B), indicating that nanomotion can be used to detect AZI susceptibility.

**Figure 2.**
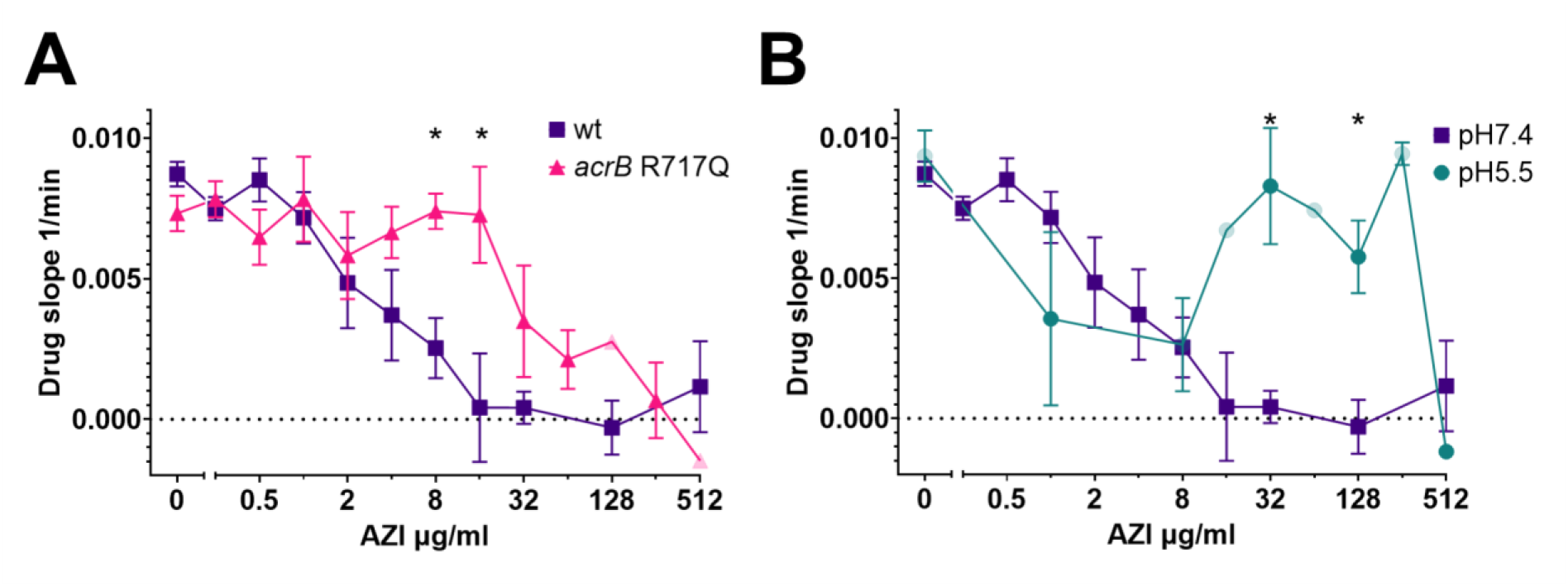
The drug-phase slope of nanomotion variance reflects the response to AZI in *Salmonella* and can be used to estimate susceptibility to the drug. A. The slope of the variance during the drug phase of the AZI-resistant *acrB* R717Q mutant and susceptible wild-type (wt) strain SL1344 at neutral pH. B. Drug slope of the wt strain at two different pH values. RT data; means ± SEM (N≥3); transparent data points shown, where N <3. *Indicates p-value <0.05 of the difference between the groups at the indicated concentration.

Drug slopes started to decline at AZI concentrations several-fold below the MIC, indicating an effect on the bacteria (Fig 2). To verify these sub-MIC effects of AZI, we used a fluorescent reporter in which the translational attenuation-based regulatory leader region (*ermCL*) is fused to GFP (Fig. 3A) instead of the native ermC methyltransferase that confers macrolide resistance (27, 28). Macrolides stall the ribosome during ErmCL translation, which opens the mRNA secondary structure allowing translation initiation of the downstream gene (28). AZI induced GFP expression in bacteria containing the reporter plasmid in a concentration-dependent manner (Fig. 3B) at these same sub-MIC concentrations where drug slopes began to decline. Maximum reporter induction was observed at or slightly above the MIC at pH 7.4, however little to no induction was seen at concentrations ≤1 µg/ml (Fig.-s 3B, S5-S6), which is in good agreement with the nanomotion data (Fig. 2). In accordance with the lower MIC at RT, the signal peaked at 4X lower concentrations at RT than it did at 37°C (Fig. 3B, S6). However, the induction levels remained significantly lower at RT, reflecting slower translation processes. At pH 5.5, GFP induction began at significantly higher concentrations compared to pH 7.4 (Fig.-s 3B, S5-S6), supporting the notion that the pH-dependence of AZI sensitivity is due to differences in antibiotic accumulation within the cell. AZI did not induce GFP at RT at pH 5.5 (Fig. 3B, S6).

**Figure 3.**
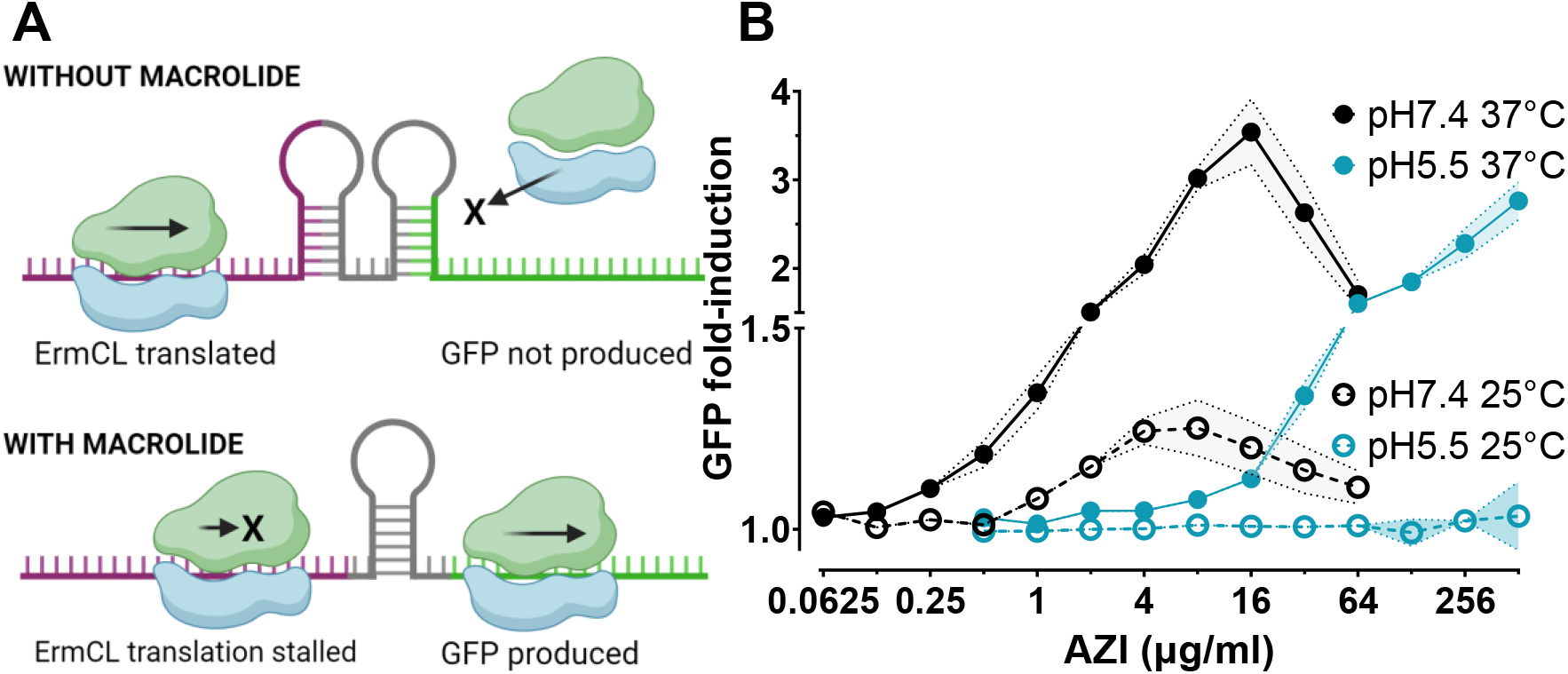
ErmCL-based reporter system was used to estimate AZI’s intracellular concentrations and effects on translation. A. Schematic representation of the reporter system. B. GFP induction of wt Salmonella after 2 h treatment with AZI. Flow cytometry data. Means ± SD (N=3).

In summary, we show that nanomotion technology can be used for rapid detection of AZI susceptibility. MIC values obtained using the standard dilution method, CFU counting results, and *ermCL*-dependent GFP induction by AZI were all consistent with the physiological responses recorded by nanomotion. Additionally, we found that nanomotion is effective for detecting PAE and assessing bactericidal activity. Our study highlights the importance of assay conditions, which significantly affected AZI efficacy and readout of the test.

## Acknowledgements

This research was funded by Estonian Research Council grants PRG335 and MOB3ERA7 Effective RApid DIagnostics and treatment of AntiMicrobial Resistant bacteria (ERADIAMR), and EU TWINNING project “Molecular Infection Biology Estonia – Research Capacity Building” (H2020-WIDESPREAD-2018-2020/GA: 857518). We are thankful to Dorota Klepacki and Alexander Mankin (University of Illinois at Chicago) for preparing and sharing the reporter. We are thankful to the whole Resistell team for helping with the nanomotion experiments.

## Author contributions

M.H. - investigation, methodology, data curation, formal analysis, visualization, writing - original draft; T.M., I.K., M.P.V. - investigation; G.C. - data curation, formal analysis, software; M.P., D.C., A.S., N.K., T.T. - conceptualization, funding acquisition, project administration, supervision. All authors contributed to the review & editing of the manuscript. Resistell AG has developed the patented (WO2023174728A1) methodology for nanomotion detection.

